# Future-proofing agrobiodiversity: climate and niche-aware conservation planning using reinforcement learning

**DOI:** 10.64898/2026.05.04.722509

**Authors:** Luca Bütikofer, Daniele Silvestro, M Luisa Rubio Teso, Ada S Molina, Carlos Lara-Romero, Raul García Valdés, Olivier Broennimann, Jose Maria Iriondo, Antoine Guisan, Blaise Petitpierre, Sylvain Aubry

**Affiliations:** Dept of Ecology and Evolution, Univ. of Lausanne, Lausanne, Switzerland; ETH Zürich, Department of Biosystems Science and Engineering, Basel, Switzerland; Swiss Institute of Bioinformatics, 4056 Basel, Switzerland; Department of Biological and Environmental Sciences and Gothenburg Global Biodiversity Centre, University of Gothenburg, Sweden; Global Change Research Institute (IICG), Rey Juan Carlos University (URJC), Madrid, Spain; Institute of Earth Surface Dynamics, Faculty of Geosciences and Environment, University of Lausanne, Switzerland; InfoFlora, Geneva, Switzerland; Swiss Federal Office for Agriculture, Bern, Switzerland; Department of Biology, Higher School of Experimental Science and Technology (ESCET), Rey Juan Carlos University (URJC), Madrid, Spain

**Keywords:** Crop Wild Relatives, Niche coverage, Climate resilience, Genetic diversity, Plant Conservation

## Abstract

Despite substantial global commitments to expand protected-area networks, the strategic allocation of limited resources remains challenging. Spatial conservation planning helps identify priority regions that maximise conservation benefits per unit area. Yet, they also tend to neglect two fundamental aspects of conservation: climate-driven range shifts and the representation of environmentally distinct populations within species. Here, we propose a continental-scale conservation planning framework that explicitly accounts for both processes through novel routines implemented in the conservation planning software CAPTAIN. We apply this framework to European crop wild relatives (CWR), for which niche coverage is a focal priority, as it underpins their potential to support agricultural adaptation to future environmental stressors through breeding programs. Comparative analyses on a subset of 186 CWR associated with five focal crops show that accounting for range shifts and niche coverage leads to substantially different conservation priorities from those obtained with a baseline model based on current distributions only. These additions reduced the number of non-protected species by 64%, increased the average protected distribution range by 43%, increased mean niche coverage from 75.8% to 84.5% and reduced the number of species with less than half of their niche protected from 35 to 10. Applied to a more comprehensive checklist of 1,140 European CWRs, the final framework identifies continental-scale priority areas representing 93.5% of these taxa and includes 94.4% of its critically endangered species. Our results highlight the importance of incorporating both temporal dynamics and within-species environmental representation when designing conservation strategies under climate change.

**Repository:** The repository will be made publicly accessible after publication at doi: https://10.5281/zenodo.19855597

## Introduction

The accelerating pace of biodiversity loss has prompted ambitious global commitments to expand protected-area networks, most notably the Kunming-Montreal Global Biodiversity Framework target to protect 30% of land and sea by 2030 (https://www.cbd.int/gbf). Yet the effectiveness of these commitments depends not merely on how much area is protected, but on where exactly protection effort is allocated (Watson et al. 2023). Spatial conservation planning (SCP) addresses this challenge by identifying configurations of protected areas that maximize conservation benefits while accounting for constraints such as limited area, economic cost, and land-use conflicts (Margules and Pressey, 2000).

Over the past two decades, SCP tools such as Zonation (Moilanen et al 2005)(Lehtomäki and Moilanen 2013), Marxan (Ball et al 2009)(Watts et al 2009), and, more recently CAPTAIN (Silvestro et al 2022, 2025) have become essential for evidence-based conservation decision-making (Moilanen et al 2009). However, current applications still often rely on simplifying assumptions that can limit their effectiveness under ongoing environmental change. In particular, species are still routinely treated as spatially uniform entities (Plumptre et al. 2025), even though populations across a species range may differ genetically and ecologically (Diniz-Filho et al. 2012), together shaping the species’ ecological niche (Pearman et al., 2010). This intraspecific variation is increasingly recognized as an important component of biodiversity because it can influence both species’ long-term persistence and their adaptive capacity (Hanson et al. 2017, Andrello et al. 2022). At the same time, this finer-scale population structure is also strongly influenced by climate change (Pauls et al., 2013), yet many conservation plans still rely on static snapshots of species distributions, even as climate change is already shifting the location and extent of suitable habitats. Although a few recent studies have begun to incorporate climate-driven range shift dynamics into SCP (Lawler et al. 2020, Chauvier-Mendes et al. 2024), particularly in aquatic ecosystems (Brito-Morales I. et al. 2022, Doxa et al. 2022), range shift dynamics are both crucial for designing resilient protected area networks, and typically omitted (Jung et al., 2024). These shortcomings are particularly consequential for crop wild relatives (CWRs), taxa that are phylogenetically close to cultivated plants (Castaneda et al., 2016). CWRs represent a particularly important target for conservation because their genetic diversity underpins crop improvement through breeding, offering potential traits for drought and high-temperature tolerance, pest and pathogen resistance, and other adaptations to environmental or biotic stresses (Dempelwolf et al., 2014). The loss of intraspecific variation for these taxa may compromise the availability of genetic diversity needed to sustain the production of food, feed, and fiber in the context of climate change (Khoury et al., 2022). Maximizing the conservation of CWRs’ ecological niche while maintaining an accessible genepool as their ranges shift under climate change should therefore be a priority. Attempts have been made to target natural populations to optimally conserve subspecific CWR diversity for single species (e.g., in wild relatives of the olive tree by Barea-Márquez et al 2026 or celery Mewis et al. 2024). However, harmonizing informative infraspecific genetic data across large and taxonomically heterogeneous species sets remains a major challenge. Environmental and geographic variables have been shown to serve as potential surrogates for intraspecific genetic variation, capturing adaptive differentiation shaped by local environmental conditions and neutral divergence driven by isolation-by-distance and isolation-by-environment (Wang & Bradburd, 2014; Hanson et al., 2017; Tobón-Niedfeldt et al., 2022). We suggest maximizing the conservation of this diversity by considering the largest possible proportion of the CWR niche, a concept we will henceforth refer to as “niche coverage”. Considering environmental variation within species in this way has already been proposed to refine both Red List assessments (Breiner et al., 2017) and protected-area networks (Hanson et al., 2020). For CWRs in particular, conservation planning that overlooks both climate-driven range shifts and the coverage of environmentally distinct populations risks missing a substantial part of the genetic variation that may be needed for future breeding and agriculture at large.

In this study, we ask how spatial conservation priorities shift when *in situ* conservation is explicitly designed to preserve both climate-driven range shifts and the niche coverage of environmentally distinct populations within species. To address this question, we implement a conservation planning framework, based on a reinforcement learning algorithm, that integrates (i) temporal range shifts and (ii) an explicit goal to maximize intraspecific niche coverage, using European crop wild relatives as a continental-scale study case (Figure 1). We compared three prioritization strategies on a subset of 186 European crop wild relatives associated with five representative and economically relevant “flagship” crops (barley, *Brassica*, lettuce, pea and wheat): a baseline model based on current distributions, a model including range shifts under climate change, and a model additionally promoting niche coverage across each species’ occupied environmental conditions. We then applied the full framework to 1,140 European crop wild relatives to derive a continental-scale prioritization plan. This step-by-step approach allowed us to evaluate the consequences of moving from static, species-level targets to a temporally dynamic framework that explicitly incorporates intraspecific diversity into *in situ* conservation prioritization. Together, these analyses allow us to assess how temporal dynamics and within-species environmental representation can reshape conservation priorities for European CWRs, and more broadly, adapt SCP to a fastly moving environment.

**Figure 1.**
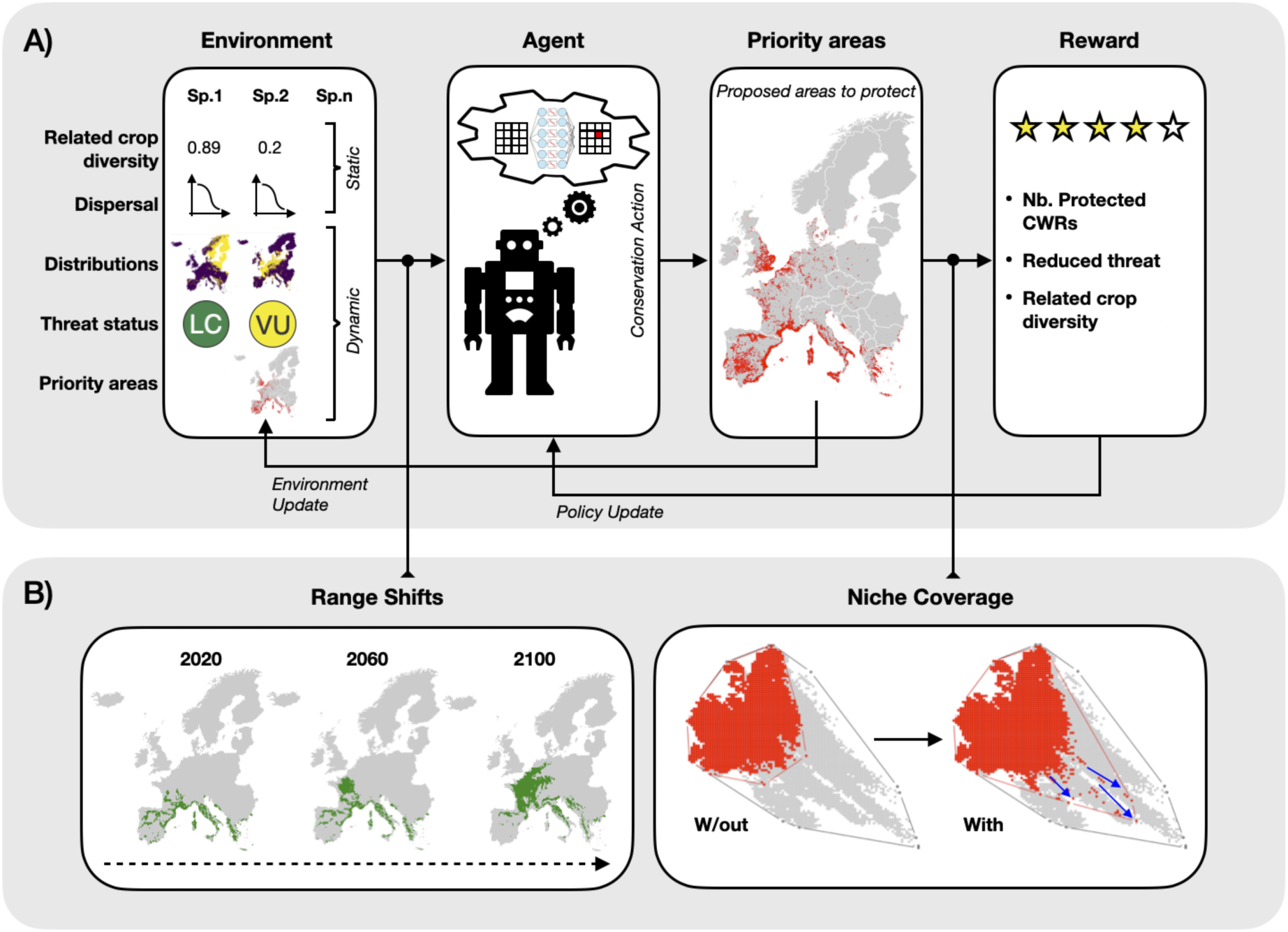
Modelling architecture to include range shifts and niche coverage in spatial conservation planning. CAPTAIN was used to compare a baseline prioritisation model (A) with a model including climate-driven range shifts and niche coverage (B). Range shifts were derived from species distribution models and simulated through CAPTAIN’s native dispersal routine, whereas niche coverage was quantified in the environmental space defined by the first two axes of a principal component analysis of 19 bioclimatic variables and incorporated through a new optimisation routine.

## Results

We compiled a list of 1,408 European native CWRs related to food and feed crops (SI *Appendix*, Table S1). The observed distribution of these European CWRs spans much of the continent, from the Southern Mediterranean islands to Southern Scandinavia (SI *Appendix*, Figure S1). However, these data showed significant observation biases in Western Europe, but were partly reduced by spatial thinning. After taxonomic harmonization and data curation, we produced species distribution models (SDMs) for 1,140 taxa. Model performance (SI *Appendix*, Figures S2, S3 and S4, Table S4) was good to excellent for all models when evaluated using AUC, and fair to good when evaluated using maximum TSS (according to Allouche et al. 2006), supporting their use in subsequent conservation-planning analyses.

Across this modeled dataset, projected habitat suitability generally declined under future climate scenarios. Median suitability decreased by 39% across all CWRs, and projected distribution centroids shifted by approximately 400 to 1,000 km (SI *Appendix* Figure S5), indicating substantial geographic redistribution of suitable conditions. This decline was stronger for taxa relatives of barley and brassicas, which showed median suitability decreases of 63% and 38%, respectively (SI *Appendix* Figure S5). Together, these results indicate that many European CWRs are expected to experience major changes in both the extent and location of suitable habitat over the coming decades.

To isolate the effect of climate-driven redistribution on conservation outcomes, we first trained CAPTAIN on the 186 CWRs associated with the five flagship crops, using predicted distribution changes under the high-emissions scenario SSP585 (O’Neill et al., 2016).

Accounting for range shifts substantially altered the resulting conservation prioritisation, with low agreement with the baseline model: only 48% of the area selected for protection by the range shifts model overlapped with the baseline model (the number of cells selected by only one model were more than double of those selected by both; SI *Appendix* Figure S6).

Accounting for range shifts, by the end of the century, added six more taxa (+3.2%) to the pool of protected and increased the average size of protected distribution range by 43% (Table 1, Figure 2B). Interestingly, these gains were not constant through time. Under present conditions, the range-shift model (27.28% mean range protection) protected 3.95% less area than the baseline model (31.13% mean range protection), but the strategy’s benefits are predicted to appear later in the century, with a protected range 9.06 percent points higher in 2050 (30.25% vs. 21.19% mean range protection for the range-shift and baseline models respectively; Figure 2E).

**Figure 2.**
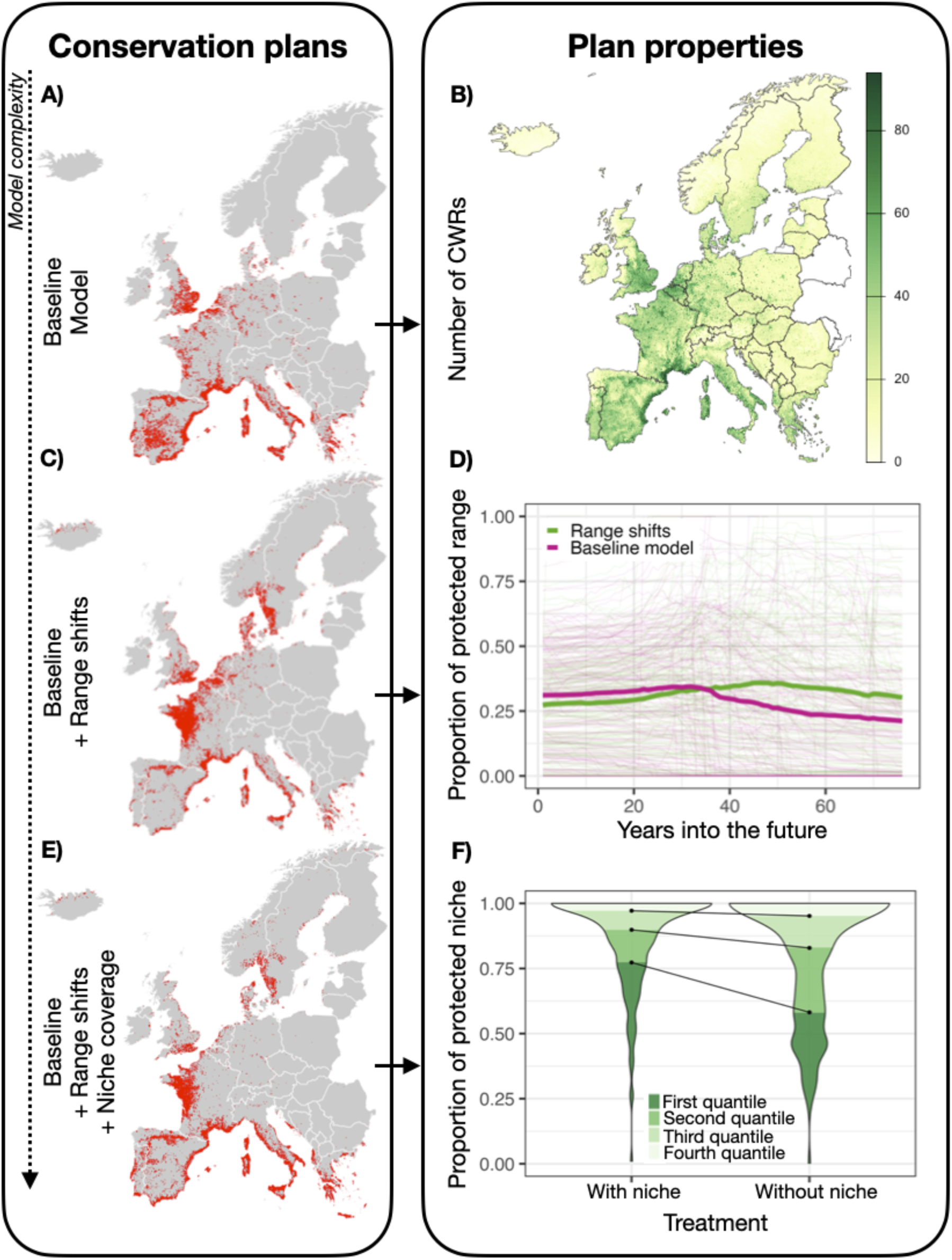
Comparison of spatial conservation plans for 186 crop wild relatives of five representative crops of economic importance: wheat, barley, pea, lettuce, and brassicas. The baseline model (A) uses potential distributions under current climatic conditions (B) to maximise the number of protected taxa while giving greater weight to threatened taxa. The range-shift model (C-D) additionally accounts for projected climate-driven redistribution of suitable habitat until the end of the century according to the high-emission scenario SSP585, based on species distribution models. The niche-coverage model (E) further incorporates the explicit maximisation of niche coverage during prioritisation, thereby increasing the proportion of each species’ niche represented within protected areas. This addition strongly reduces the number of species for which less than half of the niche is protected, as shown by the marked improvement in the first and second quartiles (F).

**Table 1.** Number of crop wild relatives (CWRs) represented by at least one protected raster cell under different conservation-planning models, shown for the present and the end of the century. Models 1–3 were evaluated on the subset of 186 CWRs associated with five flagship crops, whereas model 4 corresponds to the full framework applied to all 1,140 modelled European CWRs distributions. Model 1 is the baseline model, model 2 includes range shifts, model 3 includes both range shifts and niche coverage, and model 4 shows the final full-species prioritisation. Counts are given by threat class and in total.

|  | Nb. of protected CWR (out of 186 for mod. 1-3; out of 1140 for mod. 4) |  |  |  |  |  |  |  |  |  |
| --- | --- | --- | --- | --- | --- | --- | --- | --- | --- | --- |
| Threat class | LC |  | NT |  | EN |  | CR |  | Tot. |  |
| Time | Pres. | Fut. | Pres. | Fut. | Pres. | Fut. | Pres. | Fut. | Pres. | Fut. |
| 1- Baseline | 147 | 134 | 3 | 3 | 2 | 0 | 33 | 33 | 185 | 170 |
| 2- Range shifts | 147 | 139 | 3 | 4 | 2 | 0 | 33 | 33 | 185 | 176 |
| 3- Niche coverage | 147 | 140 | 3 | 4 | 2 | 2 | 33 | 33 | 185 | 177 |
| 4- All Spp. | 942 | 886 | 17 | 16 | 6 | 5 | 158 | 151 | 1132 | 1066 |

We assessed the effect of adding niche coverage to the range-shift model using the subset of 186 CWRs associated with five flagship crops. The niche coverage model produced a conservation prioritisation map characterised by a moderate agreement with the range shifts model: 64% of the area selected for protection by the niche coverage model overlapped with those selected by the range shifts model (SI *Appendix* Figure S6). Including this component increased the mean proportion across each species’ protected niche from 75.8% to 84.5%, with 155 out of 186 species reporting an increase in their proportion of protected niche. In particular, under the model accounting for range shifts alone, 35 taxa had less than 50% of their niche protected, whereas this number fell down to 10 taxa when niche coverage was included.

We then applied the full conservation-planning framework, including both range shifts and niche coverage, to the complete set of 1,140 European crop wild relatives. The resulting continental-scale prioritization identified a distinct spatial pattern of conservation importance, with major priority areas located in southern Scandinavia, along the Atlantic western coast, and across large parts of the Mediterranean shores (Figure 3). These areas therefore, emerged as the main candidates for long-term *in situ* conservation when both future climatic redistribution and within-species environmental representation are considered simultaneously. This conservation prioritisation plan covers 93.5% of all CWRs (1066 taxa), including 94.4% of CR, 83.3% of EN, 88.9% VU taxa (Table 1).

**Figure 3.**
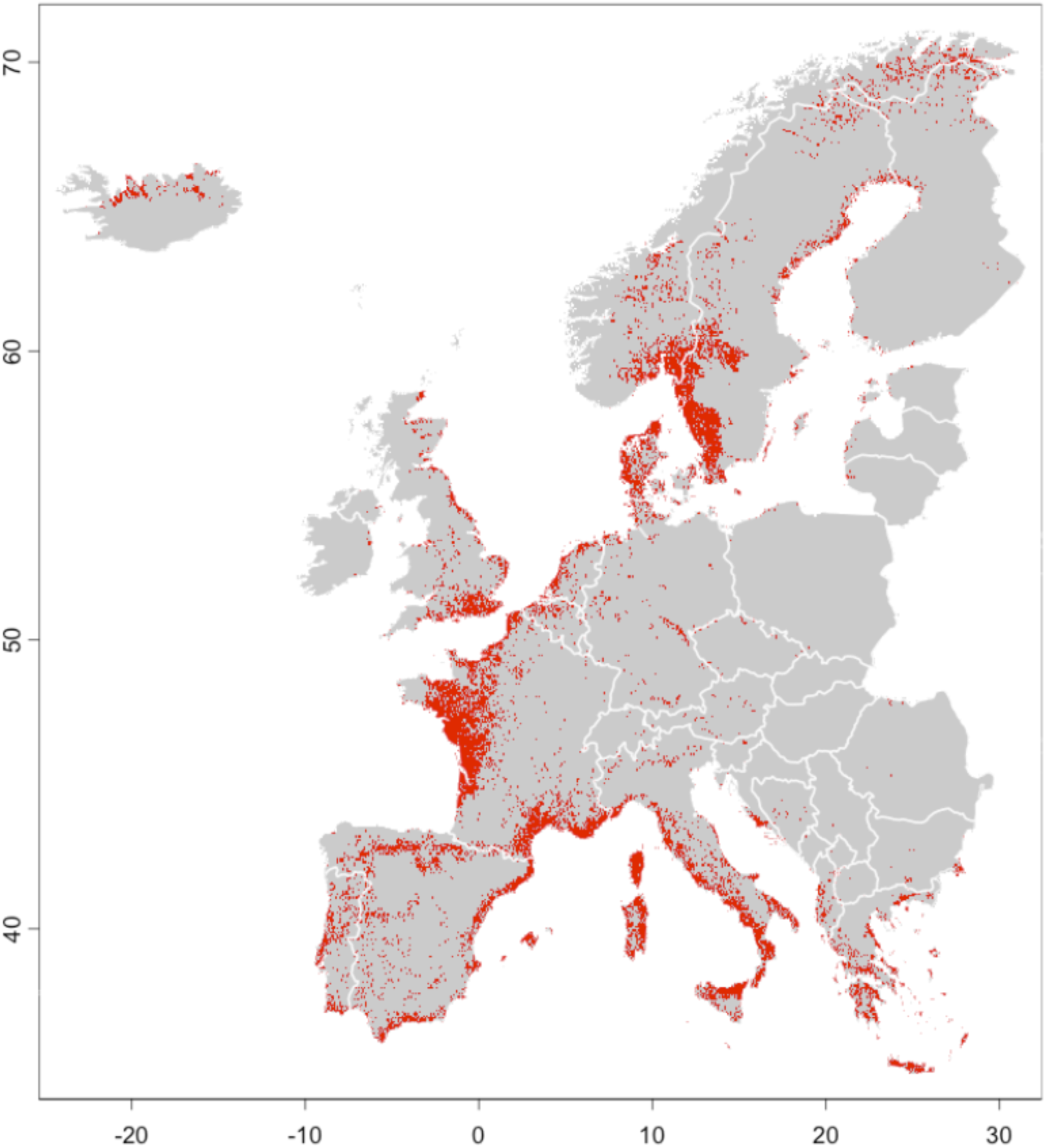
Spatial conservation plan for European crop wild relatives (CWR) accounting for 1,140 taxa and their range shifts until the end of the century according to the high-emission scenario SSP585. The prioritisation was designed to maximise the number of protected CWR taxa, while prioritising threatened species, ensuring balanced representation of their related crops, and increasing the proportion of each species’ represented within the priority area.

Because training conservation-planning models at a continental scale is computationally demanding, we also evaluated whether an SCP model trained on a subset of the 186 CWRs could predict the priorisation patterns for a broader set of taxa. Our generalisation test showed good performance when CAPTAIN was trained on flagship and projected to the 1,140 CWRs. 62% of the area selected for protection by the model trained with all species overlapped with the area selected by the model trained with flagship species only. The cells selected by only one model, however, were very sparse; aggregating the overlap map at 15 x 15 km^2^ resolution (Figure S7), showed that most disagreement areas only contained very few pixels selected for protection (6.8% on average). When testing the value of the flagship CWRs as umbrella species, we found that the largest priority areas identified by CAPTAIN were the same for both models (*i.e.* accounting for all European CWRs *vs* including flagship CWRs only; Figure 2). However, the two conservation plans diverged in some areas (Figure S7). 59% of the area selected for protection by the model trained with all species overlapped with those selected by the model trained with flagship species only. Although this result is not very different from the generalisation comparison counterpart, the cells selected by only one model were less sparse. Indeed, aggregating the overlap map at 15 x 15 km^2^ resolution, showed that most disagreement areas contained several pixels selected for protection (50% on average).

## Discussion

Accounting for both climate-change-driven range shifts and niche coverage, increased both the number of CWRs represented in conservation priority areas and the average proportion of protected niche (Figure 3 & 4). Interestingly, this does not come at the cost of reducing the number of protected taxa—a result that was not guaranteed a priori, given that both mechanisms primarily reshape where protection is allocated rather than how many species fall within protected areas (Levin et al. 2015). Conservation plans are often evaluated essentially on the number of species covered by the areas selected for conservation (e.g. Vincent et al., 2019 for CWR) and therefore may substantially underperform in preserving the intraspecific diversity and long-term climatic persistence that are critical for effective conservation (Jung et al. 2024, Giakoumi et al. 2025; Tobón-Niedfeldt et al., 2022). This is especially important in the context of agrobiodiversity, much of which is threatened by genetic erosion that requires urgent conservation measures (Khoury et al., 2022). In contrast to cultivars or landraces that are often conserved *ex situ*, in genebanks or seed repositories, CWR are mostly wild populations that require dedicated conservation measures, referred to as *in situ* conservation (Jago et al., 2024). Most monitoring attempts to prioritize *in situ* conservation of CWR so far have been restricted to the actual distribution (SI *Appendix* Table S1) with relatively few cases where climate has been considered (but see Van Treuren et al. 2020; Phillips et al. 2017; Aguirre-Gutiérrez et al. 2017; Rahman et al. 2023). Conservation planning that ignores these dynamics risks investing in areas whose conservation value will erode over the coming decades while neglecting regions that will become critical in the future (Jung et al., 2024).

The spatial priority areas identified by our model for all European CWRs reveal a temporal shift in the geography of CWR conservation (Figure 3). Under current conditions, the Mediterranean coast emerges as the primary area of importance, consistent with its role as a center of CWR diversity and endemism (Vincent et al., 2022). However, as climates shift, the North Sea coast gains priority as the taxa currently occupying the Mediterranean shores will track their climatic niche till the North Sea, a northward shift consistent with previous studies in this area (Aguirre-Gutiérrez et al. 2017). This temporal shift also helps explain a pattern that might initially appear counterintuitive: range-shift and niche-coverage models assign substantial priority to southern Europe, where local extinctions due to climate change are most likely (Figure 3). This reflects that our model optimizes protection over the full period, from the coming decades to the end of the century, rather than at a single future time point. Protecting CWRs in these southern locations in the near term buys critical time for assisted migration, natural dispersal into newly suitable areas, and targeted *ex situ* collection of locally adapted populations (Butt and Gallagher 2018; Simonson et al. 2021). This highlights the importance of viewing in situ conservation as a temporally dynamic strategy rather than a permanent commitment (Reside, Butt & Adams 2018, McLaughlin et al. 2022).

Within the broader set of available spatial conservation planning software, Zonation (Moilanen et al., 2022) can approximate temporal range-shift dynamics using distribution maps across different time steps and connectivity layers (Honeck et al., 2020). This is a useful and computationally cheaper proxy. However, it differs from CAPTAIN’s explicit dispersal simulation in one important respect: CAPTAIN’s temporal framework allows early conservation action to alter a taxon’s threat status, which in turn changes its future conservation value and influences subsequent planning decisions—a feedback loop that static time-step approaches cannot replicate. Similarly, while environmental variables offer a promising and cost-effective proxy for intraspecific genetic diversity in conservation planning (Hanson et al., 2017), their effectiveness is not universal: surrogate-based prioritisations have been shown to perform inconsistently across amphibian and reptile species—in some cases no better than random selection—(Hanson et al., 2021), underscoring the value of integrating direct genetic data when available.

A practically important finding emerges from our efficiency tests: CAPTAIN models trained on flagship CWRs generalized well to the full set of European crop wild relatives, with an average spatial discrepancy of only 7% and broadly consistent identification of the largest priority areas. Although this approximation does not fully replace training on the complete dataset, it suggests that reinforcement-learning-based conservation planning (Silvestro et al. 2022) can be applied at reduced computational cost while retaining much of the resulting prioritisation pattern, a practically important finding given the energy and time demands of model training (Gardner *et al*., 2025).

Restricting the model training on a subset of the entire CWR checklist can also be justified from a pragmatic point of view, as representing some of the most economically valuable subset of genetic resources for food and agriculture. More generally, the use of a subset of taxa as conservation surrogates or umbrella species has long been proposed as a way to simplify spatial conservation planning (Caro and O’Doherty 1999, Roberge and Per Angelstam 2004), but its effectiveness remains context dependent and often limited (Branton and Richardson 2011; Tälle et al. 2023). In our case, prioritization trained on the flagship subset and projected to the full set of CWRs captured broad spatial patterns but still resulted in losses relative to prioritisation trained directly on the full CWR dataset. There is a fine line between prioritizing globally to provide coherent, politically and economically workable conservation planning and accounting for ecological complexity and dynamics. A more comprehensive coverage, possibly also integrating finer phylogenetic distances to crops as well as functional diversity, may yield a further improvement in defining conservation targets (Chauvier-Mendes et al., 2024).

We have shown that accounting for niche coverage and climate-driven range shifts can improve conservation planning at a continental scale. However, these results should be interpreted in light of several limitations. First, the occurrence data used to build species distribution models may be subject to observational bias (Beck et al., 2014). Here, despite applying spatial thinning (Boria et al., 2014), a bias towards Western and Northern Europe is evident and may consequently result in lower prioritization (due to lower sampling). Second, our approach does not account for neonative CWR populations originating outside the study area, particularly in Eastern Europe and Turkey, from where some migration may be underestimated. Third, we applied a single dispersal kernel across all species when simulating range shifts. While this represents a substantial improvement over neglecting dispersal entirely, species-specific parameterization would yield more accurate projections where data on dispersal capacity are available. A possible improvement to this approach would take into account species’ dispersal biology and use a different kernel for each taxon or taxon group. Finally, our species distribution models carry the intrinsic limitations of the correlative approach, including the omission of biotic interactions and the assumption that species do not adapt to novel climatic conditions.

As protected-area networks are expanded to meet the targets of the Kunming-Montreal Global Biodiversity Framework, the question of where to place protection is inseparable from the question of what, exactly, we are trying to protect. If the goal is merely to record species presence within protected boundaries, static approaches to conservation planning may suffice. But if the goal is to safeguard intraspecific diversity—*e.g.*, locally adapted populations, climatically marginal ecotypes, and the genetic variation that breeders will need to develop crops capable of withstanding conditions that do not yet exist (Cortes et al., 2021), then conservation planning must become both more dynamic and more ecologically precise. The approach developed here, integrating explicit range-shift dynamics and niche-coverage optimization into a reinforcement-learning framework, represents a step in that direction. It will not be the last: as species-specific dispersal data accumulate (Lososová et al. 2023), as genetic resources become better characterized (Khoury et al., 2022), and as climate projections improve (Brun et al. 2022), conservation planning tools will need to evolve alongside them. In a world where the genetic diversity of crop wild relatives may determine the resilience of our food systems, that is an argument for not merely protecting more land, but also for ensuring that what we protect today will still be worth protecting tomorrow.

## Methods

### Species distribution models

We compiled an initial checklist of 1,408 food and feed CWR taxa for Europe from multiple CWR literature sources (Appendix SI Table S1), excluding non-native taxa. We retrieved global occurrence data for these taxa from GBIF and Genesys APIs (https://GBIF.org; https://www.genesys-pgr.org; https://doi.org/10.15468/dl.egyw3p; https://doi.org/10.15468/dl.6r3qp2; https://doi.org/10.15468/dl.gthgux). Data were downloaded using the R environment (R Core Team 2023) with ‘rGBIF’ (Chamberlain and Boettiger, 2017) and ‘genesysr’ (Obreza, 2019) packages. Raw occurrence data were subsequently curated for taxonomic harmonisation and geographic accuracy, discarding points with less than two kilometres of accuracy or imprecise location (Rubio Teso et al., 2020, 2022). To reduce model bias towards more densely sampled environments, we thinned each species’ occurrences spatially so that no two points were closer than 2 km. Furthermore, if multiple occurrences occurred at locations with the same values for all predictors, only one was retained, thereby removing duplicates in environmental space.

To save computation time and reduce resource consumption during model training, all species with more than 10,000 occurrences were further spatially thinned by increasing distance thresholds in 1 km steps until the number of resulting occurrences was reduced to 10,000 or fewer. We sampled pseudoabsences as 10,000 random points worldwide stratified along climatic gradients.

We used a diverse set of predictors covering climate, topography, soil and land-use (SI, SDM predictors). We used future climate data from two global circulation models (GCMs) and two Shared Socioeconomic Pathways (SSPs): MPI-ESM1.2-HR (Mauritsen et al., 2019) and UKESM1-0-LL (Good et al., 2019), because they represent extreme ends of low (MPI-ESM) and high (UKESM) climate sensitivity (Lange, 2021), and SSP 370 (moderate) and 585 (extreme) to cover a wide range of forcing scenarios.

We used three nested tiers of model sophistication based on the number of occurrences available (SI *Appendix* Tables S2 & S3). First nested tier (1), the “Principal Component Model (PCM)”, targeted CWRs with more than 5 occurrences (1140 taxa) were modelled with a low sophistication approach based on the density of occurrences in the 2D surface formed by the first two components of a PCA (Broennimann et al. 2012) carried out using 19 bioclimatic variables. The second nested tier (2), referred to as “Global”, comprises CWRs with more than 30 occurrences globally (*i.e.* at world scale; 916 taxa), which were modelled with an ensemble of three algorithms using the 19 bioclimatic variables. We will refer to this level of model sophistication as (3) Species with more than 30 occurrences in Europe (826 taxa) were modelled at this continental scale with an ensemble of three algorithms based on the prediction of the Global model as an input covariate (Guisan et al. 2025), additional bioclimatic variables, soil, topography, and land-use/land-cover variables. We will refer to this level of model sophistication as “Regional”.

### Spatial conservation planning

We identified the most important areas for the conservation of CWRs in Europe with the CAPTAIN v.2 software (Silvestro et al., 2022). CAPTAIN implements a spatially-explicit simulation of biodiversity (the *environment*) that evolves over time and within which its *agent* selects areas for protection. It uses reinforcement learning to optimise the behaviour of the agent and maximise a reward, e.g. quantifying biodiversity outcomes of conservation.

The environment is a stack of SDMs for each target taxa, and is temporally dynamic as species track their climatic niche through a diffusion process. At each time step, the agent observes the current state of the environment and chooses a number of cells for protection based on rules—the model’s *policy*—which are initialised randomly, and refined during training. A reward is computed based on the outcome of the agent’s policy, and accumulated over time. At the end of the training, the agent has learned the conservation policy. Training aims to optimize the behavior of the agent (its policy) to maximize the total reward, which is based on the number of taxa protected, their degree of threat according to the IUCN red list classification, the number of related crops, and the novel reward to maximise niche coverage (Supplementary Methods, Reward definitions and policy network). To test for the effects of range shift and the niche coverage, we performed three runs on the flagship CWRs: one without accounting for range shifts or niche coverage (the “baseline” model), one accounting for Range Shifts (the “RS” model), and one accounting for both Range Shifts and Niche Coverage (the “RSNC” model). To quantify the effects of niche coverage, we created a new indicator of niche protection as follows. We first derived the number of one km^2^ cells of the environmental plane on which the species is present, and the number of cells of the environmental plane on which the species is present and protected. We then derived the Minimum Convex Polygons (MCP; REF) of both the presence and protected cells. The niche protection indicator was then the ratio of the areas of the protected MCP over the presence MCP.

To test the usage of flagship CWRs as an umbrella subset (McGowan et al., 2020), we trained the RS model both on the flagship only and on all CWRs. Subsequently, to test the generalisation capabilities of CAPTAIN, we used the RS model trained on the flagship CWRs to generate a conservation plan for all the CWRs. Finally, we compared the outputs for these two pairs of conservation exercises by computing the spatial difference between each pair (e.g., the flagship RS model minus all CWR RS model). This resulted in maps showing areas selected by both runs, areas selected by neither, and areas selected by only one (SI *Appendix* Figure S7).

The software routines developed for this study, enabling both range-shift dynamics and niche coverage optimisation, are freely available as part of the latest CAPTAIN release (https://www.captain-project.net).

## Author contributions

Luca Bütikofer: Conceptualization, Data curation, Formal analysis, Investigation, Methodology, Software, Validation, Visualization, Writing - Original Draft, Writing - Review & Editing;

Daniele Silvestro: Conceptualization, Formal analysis, Investigation, Methodology, Software, Validation, Writing - Original Draft, Writing - Review & Editing;

Luisa Rubio Teso : Conceptualization, Data curation, Formal analysis, Funding acquisition, Investigation, Writing - Original Draft, Writing - Review & Editing;

Ada Molina: Conceptualization, Writing - Original Draft, Writing - Review & Editing;

Carlos Lara-Romero : Conceptualization, Data curation, Formal analysis, Investigation, Writing - Original Draft, Writing - Review & Editing;

Raul García Valdés : Conceptualization, Writing - Original Draft, Writing - Review & Editing;

Olivier Broennimann: Conceptualization, Methodology, Resources, Visualization, Writing - Original Draft, Writing - Review & Editing;

Jose Maria Iriondo : Conceptualization, Formal analysis, Funding acquisition, Investigation, Project administration, Resources, Supervision, Validation, Writing - Original Draft, Writing - Review & Editing;

Antoine Guisan: Conceptualization, Methodology, Project administration, Resources, Supervision, Validation, Writing - Original Draft, Writing - Review & Editing;

Blaise Petitpierre: Conceptualization, Data curation, Formal analysis, Funding acquisition, Investigation, Methodology, Project administration, Supervision, Validation, Writing - Original Draft, Writing - Review & Editing;

Sylvain Aubry: Conceptualization, Funding acquisition, Investigation, Methodology, Project administration, Supervision, Validation, Writing - Original Draft, Writing - Review & Editing;

## Acknowledgement

The research has been supported by the Horizon project COUSIN funded by the EU (European Union), the Swiss State Secretariat for Research and Innovation (grant n°22.0412). D.S. received funding from ETH Zurich and the Swedish Foundation for Strategic Environmental Research MISTRA within the framework of the research programme BIOPATH (F 2022/1448).

## Appendix

### Supplementary Methods

#### SDM predictors

##### Global climate

Global climate data was obtained from Chelsa V 2.1 (Karger et al. 2017, Karger et al. 2021). We selected the variables in the BIOCLIM set.

##### Terrain

Slope and terrain roughness indicators were computed from Copernicus’ 25 m x 25 m Digital Elevation Model (EU-DEM, not maintained anymore and no longer distributed by Copernicus) with the “terrain” function from the package “terra” under R (V 4.2.2) using eight neighbouring cells and subsequently aggregating the results at 100 m x 100 m resolution. Namely, we computed TRI (Terrain Ruggedness Index), the mean of the absolute differences between the value of a cell and its 8 surrounding cells, TPI (Topographic Position Index), the difference between the value of a cell and the mean value of its 8 surrounding cells, and “Roughness”, the difference between the maximum and the minimum value of a cell and its 8 surrounding cells. Slope was computed according to the Horn method (Horn, 1981).

##### Land use

Land-use/land-cover (LULC) variables for Europe were derived from the CORINE Land Cover raster from Copernicus (version 18, 100 m x 100 m resolution, CORINE, 2019). Original raster files were one-hot-encoded (Hancock and Khoshgoftaar, 2020), producing one raster file per land-use class, indicating for each pixel the presence or absence of that class. Moving-window analyses with various window sizes (square windows with edge lengths of 500, 1100, 2600, and 5100 m) were performed on the one-hot-encoded files. Both operations were carried out using whitebox tools V 2.4.0 (Lindsay, 2014).

##### Soil

Chemico-physical soil properties were obtained from ISRIC—World Soil Information’s SoilGrids (Poggio et al., 2021) for the soil depths of 0 to 5 cm and 5 to 15 cm.

##### Wetness index

The topographic wetness index was computed with Whitebox Tools’ WetnessIndex function (Lindsay, 2014). This index maps “the propensity for a site to be saturated to the surface given its contributing area and local slope characteristics” (WhiteboxTools User Manual). The input DEM was a 100 m x 100 m resolution DEM derived through spatial aggregation of the 25 m x 25 m resolution Copernicus (mean of 25 m x 25 m values within each 100 m x 100 m pixel).

##### Solar radiation

Potential, incoming, direct and diffuse solar radiation [kJ / m²] for each month of 2024 (thus with a 28-day February) was computed with the default method “Height of Atmosphere and Vapour Pressure” in SAGA-GIS “Potential Incoming Solar Radiation” module ( https://saga-gis.sourceforge.io/saga_tool_doc/2.1.4/ta_lighting_2.html). The default atmospheric height of 12000 metres and the default vapour pressure of 10 mbar were used. Solar radiation was computed for each month accounting for topographic shading; topography was provided with a 100 m x 100 m resolution DEM derived through spatial aggregation from the 25 m x 25 m resolution Copernicus (mean of 25 m x 20 m values within each 100 m x 100 m pixel).

To account for the canopy filtering in forests, the direct (1) and diffuse (2) components of the potential incoming solar radiation were treated differently. (1) Direct radiation for cells that correspond to Corine forest land-cover classes (23, Broad-leaved forest; 24, Coniferous forest; 25, Mixed forest) was reduced according to Beer’s Law “radiation below = radiation above * transmission coefficient”, where “transmission coefficient = exp(-K*LAI)” (ref. 4). Monthly “LAI” (Leaf Area Index) data from June 2018 to April 2019 were sourced from Copernicus (since June 2018 and May 2019 were missing from the dataset, we used May 2018 and April 2019 as substitutes) and disaggregated to 100 m x 100 m resolution. Gaps in the disaggregated LAI layers due to cloud cover were filled with an inverse-distance weighted algorithm using a search radius of 2100 m and a weight (i.e. the power parameter) of 1 (function “FillMissingData” from Whitebox Tools). “K” (extinction coefficient, Equation 1) was computed according to the formula below (ref. 5) at hourly time-steps for each month according to the hourly solar angle “θ” (computed with “solalt” function from the “microclima” (version 0.1.0, https://rdrr.io/github/ilyamaclean/microclima/) R package), and typical “X” values (i.e. leaf angles distribution, the ratio of vertical to horizontal projections of leaf foliage) for the three Corine forest classes (1.2, Broad-leaved forest; 0.4, Coniferous forest; 0.8, Mixed forest) sourced from the “laifromhabitat” function of R package “microclima”. To reduce computation time, hourly values for the extinction and transmission coefficients were computed every two hours and every six days. Monthly values were derived from these hourly data by averaging (*i.e.* mean).

(2) Canopy effects on diffuse radiation for cells that correspond to Corine forest land-cover classes were computed with the same law used for the direct component (Beer’s law), but without “K” since diffuse radiation is isotropic and is not affected by the leaf angle distribution.

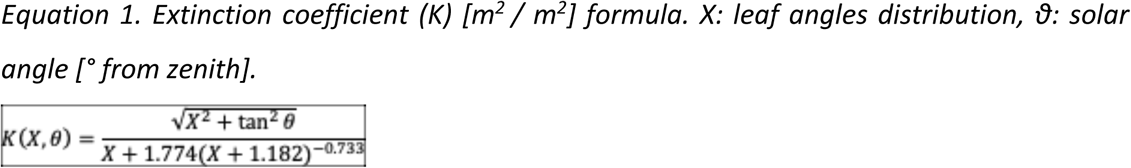

Monthly values for direct and diffuse radiation were summed to produce monthly estimates of the potential incoming radiation. Monthly potential incoming radiation values were then summed to derive yearly values.

##### Species distribution modelling

To compensate for an imbalance in the number of occurrences vs. background points, we downsampled the background class to match the number of points in the occurrence class. However, to ensure sufficient coverage of the background environment, we required at least 500 background points. To compensate for class imbalance in species with fewer than 500 occurrences, we used the inverse of class prevalence as weights (e.g., occurrences were weighted twice as heavily as background points for species with 250 occurrences and 500 background points).

We modelled each species with each algorithm for all combinations of GCMs, SSPs, and time-steps. Then, for each combination of species, SSPs and time-steps, we averaged the results of the three algorithms for the two GCMs—resulting in one potential distribution map for each species*time-step*SSP combination (*i.e.* six maps per species). This average was weighted by the algorithm’s performance, as indicated by the True Skill Statistic (TSS), computed for the binarisation threshold that maximises TSS (*i.e*., max.TSS); the two GCMs were given equal weights, effectively averaging away their differences. Outputs based on different GCMs from PCA models—for which no ensembling was performed—were averaged using the unweighted mean.

For each time-step*GCM*SSP combination, we performed MOP (mobility-oriented parity, Owens et al., 2013) analysis to detect novel climate areas, to which we assigned a presence probability of zero, since occurrence and background points were sampled within the species’ ecoregion, there is no reason to expect the species to occur in areas of novel climate.

##### SDM performance

Model performance was evaluated with the maximum True Skill Statistic (max TSS), Area Under the Curve (AUC) of the Receiver Operating Characteristic (ROC) curve, Correlation, and maximum Kappa (Figure SI2). These metrics were computed for all model sophistications, but only for species with more than 30 occurrences were obtained from cross-validated data; this means that all algorithms can be compared based on cross-validated data, even though PC models are effectively used in the conservation planning only for species with less than 30 occurrences, for which cross-validation is not possible. Cross-validation was performed by 10-fold (GLM and RF) and 5-fold (SVM, to speed up computations for the slower SVM function) splits of the training data. Cross-validated model predictions were used to compute performance metrics of the individual algorithms. They were also averaged using a weighted mean, with the same max-TSS-based weights as for the ensembling procedure, to cross-validate the ensemble models.

##### CWR habitat suitability trends and range shifts

We derived a measure of habitat suitability trend by computing the difference between future (2071-2100 for ssp370 and ssp585) and present suitability values for each pixel. We then summed the difference values for each CWR. Since northern pixels in the WGS84 reference system are larger, we weighed the difference value according to the pixel actual surface area. We computed range shift distance estimates by deriving the distance between each CWR current distribution centroid and that of the future distribution (2071-2100 for ssp370 and ssp585).

##### CWR diversity and turnover

We computed indicators of species diversity and turnover for all CWRs in our list and for all CWRs of each of the flagship crops. We binarised our SDMs into potential presences (1) and absences (0) using the threshold that maximises the TSS. Then, we computed the number of species as the sum of the binarised rasters across time steps and emission scenarios. We also calculated turnover metrics for transitions between the present and the two future time-steps for each emission scenario (*i.e.*, from the present to 2041-2070 for ssp370 and ssp585, and to 2071-2100 for ssp370 and ssp585). We computed the potential numbers of extinct species (L), persistent species (P) and gained neonative species (G, *i.e.*, future-suitable species).

##### Nearest crop

Phylogenetic data were derived from the most recent megatree included in GIFT (Zuntini et al., 2024). The related crop for each CWR of the European list was identified by pruning away from the tree the terminal taxa not included in the list or FAO’s crop list, and selecting for each CWR the nearest crop in terms of cumulative, connecting branch length. The related crops were only used whenever there was no related crop listed in the source publication for that CWR, or there was no single crop sharing the same genus.

##### Threat status

We used the SDM projections for the current climate to derive a proxy for the threat class of the 600 CWRs for which no IUCN red list assessment is available. As a proxy for threat status in the absence of an IUCN assessment, we computed the Extent of Occurrence (EOO) and Area of Occupancy (AOO) for each CWR based on our binarised SDMs (land-use filtered) using the R package red (v 1.6.2). A species is classified as Critically Endangered (CR) if its EOO is less than 100 km² or its AOO is less than 10 km². If its EOO is under 5,000 km² or AOO under 500 km², it is classified as Endangered (EN). If its EOO is under 20,000 km² or AOO under 2,000 km², it is classified Vulnerable (VU). Species exceeding all these thresholds are classified as Least Concern (LC).

To test the accuracy of the computed Criterion B-based IUCN threat status, we compared modelled values to empirical values for all species with an official assessment. To this end, we assigned an integer representing the threat rank to each threat class (*i.e.* LC=1, NT=2, VU=3, EN=4, CR=5, EW=6, EX=7; DD and NE were excluded) and computed the difference (*i.e.* assessed minus computed; therefore, negative numbers mean overestimated threat, and positive numbers underestimate threat).

When comparing the taxa with an assessed threat class with their computed threat class, most computed threat classes were the same as the assessed ones (i.e., 352 taxa), for 67 taxa the threat status was overestimated, and 14 were underestimated (SI, Figure S10). Note that the number of taxa with both assessed threat status and SDMs was only 433, since some of the assessed CWRs did not have enough occurrences with which to produce SDMs.

##### Reward definitions and policy network

We defined three reward signals that captured different aspects of conservation outcomes, namely the protection of the CWR range, the equal protection of the related crops, and the protection of their environmental niche (Figure 1). The first reward was computed, at each time step, as the average fraction of protected range across species, weighted by their extinction risk status (Silvestro et al., 2025 A&B). This measure changed over time as a function of: the increasing number of protected cells implemented by the agent, the shifting range of species as they tracked their moving environmental niche, their simulated changes in extinction risks based on the predicted expansion or contraction of their range. The second reward was based on the average fraction of protected range across species weighted by the inverse of the number of CWRs related to the same crop, therefore prioritising equal crop representation in the protected area. The third reward quantified the proportion of protected niche. The niche is here defined as the inhabited range of climates derived from binarised SDMs, and is computed from the first two principal components of the 19 bioclimatic variables used in the SDM training. For each CWR, we computed the minimum spanning tree of the graph formed by plotting all protected pixels for that CWR on the 2D surface formed by the first two principal components of the 19 bioclimatic variables —we will henceforth refer to this surface as the “environmental plane”. The niche coverage reward was set to be proportional to the total length of this minimum spanning tree.

The policy network was a fully connected feedforward neural network with two hidden layers of 8 and 2 nodes and ReLU activation functions. The input of the network included, for each cell, a set of features computed at each time step based on the current state of the environment. The features included the number of species in each extinction risk class and the same numbers computed based on future habitat suitability. They also included the number of species in each extinction risk class after excluding species that are already included in protected areas. Finally, we included a feature quantifying, for each cell, the minimum environmental distance between each non-protected cell and the closest protected cell. This distance was computed as the Euclidean distance on the environmental plane. The output of the network was a score from which the top 10 highest ranking cells were selected for protection before recomputing the features. The agent’s space of action was set to identify 1560 protected cells (10% of the study area) selected over 35 time steps (years), while the environment was evolved for a total of 75 time steps, with the reward cumulated over the entire time frame.

We trained the models using CAPTAIN’s parallelized evolution strategies (a genetic strategy algorithm; Silvestro et al., 2025 B, Salimans et al., 2017, Sutton et al., 1999) to estimate the weights of the policy network that maximize the expected reward. We used nine parallel environments based on the nine decimated datasets and trained the main model over 1000 epochs. We first trained the range shifts model, and then fine-tuned the additional models: the baseline model by removing range shifts, and the niche model by activating the niche coverage routine.

## Supplementary figures

**Figure S1:**
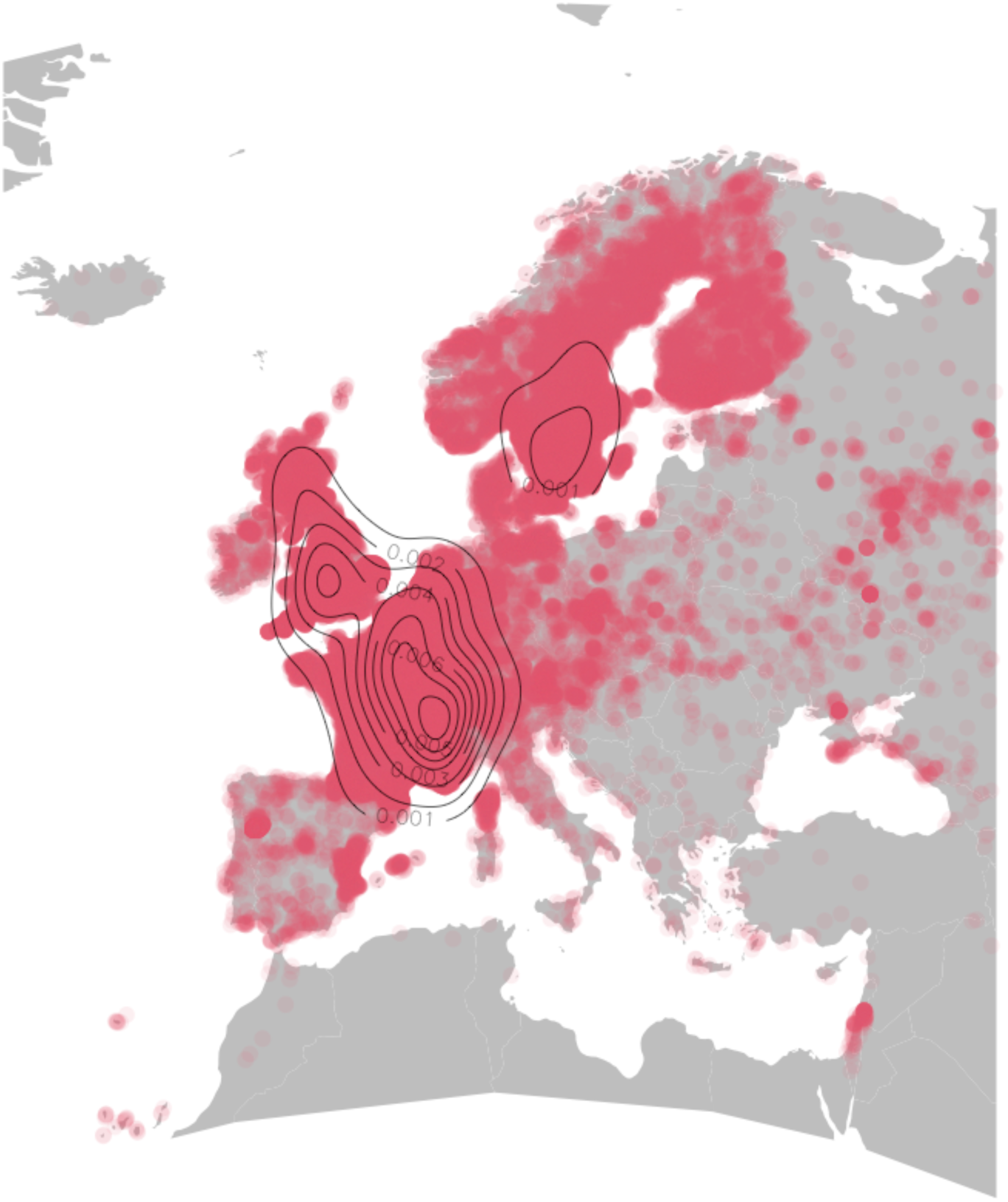
Occurrence points density for all European CWRs (kernel density estimates overlaid as isolines). Note the observation bias in the distribution of occurrences, with few points in southern and eastern Europe.

**Figure S2.**
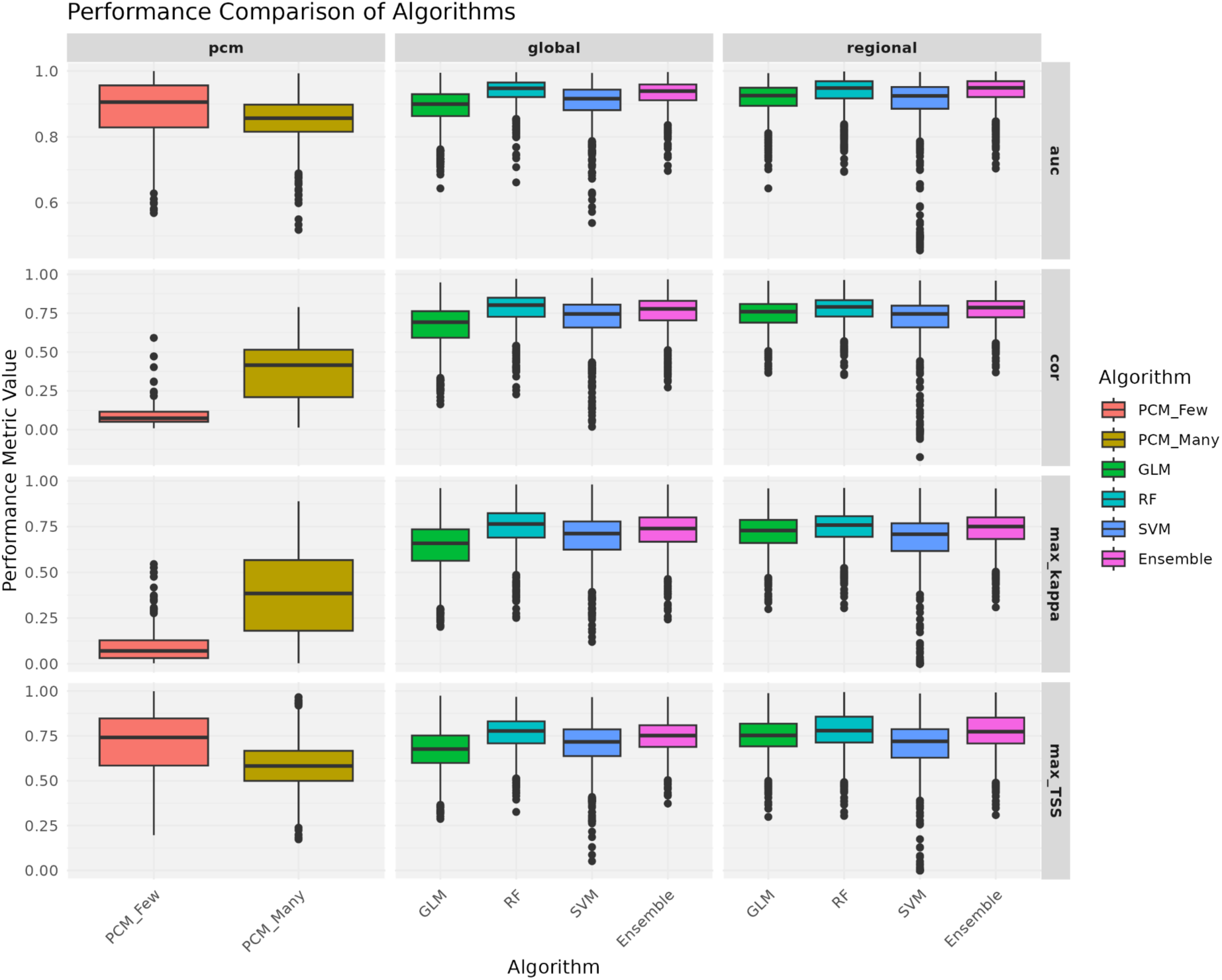
SDM performance indicators. Each row of boxplots corresponds to a performance indicator: Area Under the Curve (AUC) of the Receiver Operating Characteristic (ROC) curve, Correlation, maximum Kappa and True Skill Statistic (max TSS) each column of boxplots corresponds to a tier of model sophistication (PCM global/many, and regional/few), and each colour corresponds to a modelling algorithm (GLM, RF, SVM, and Ensemble of the previous algorithms). Note that the PCM_Few is not cross-validated due to the low number of occurrences. Scoring performances in Table S4

**Figure S3.**
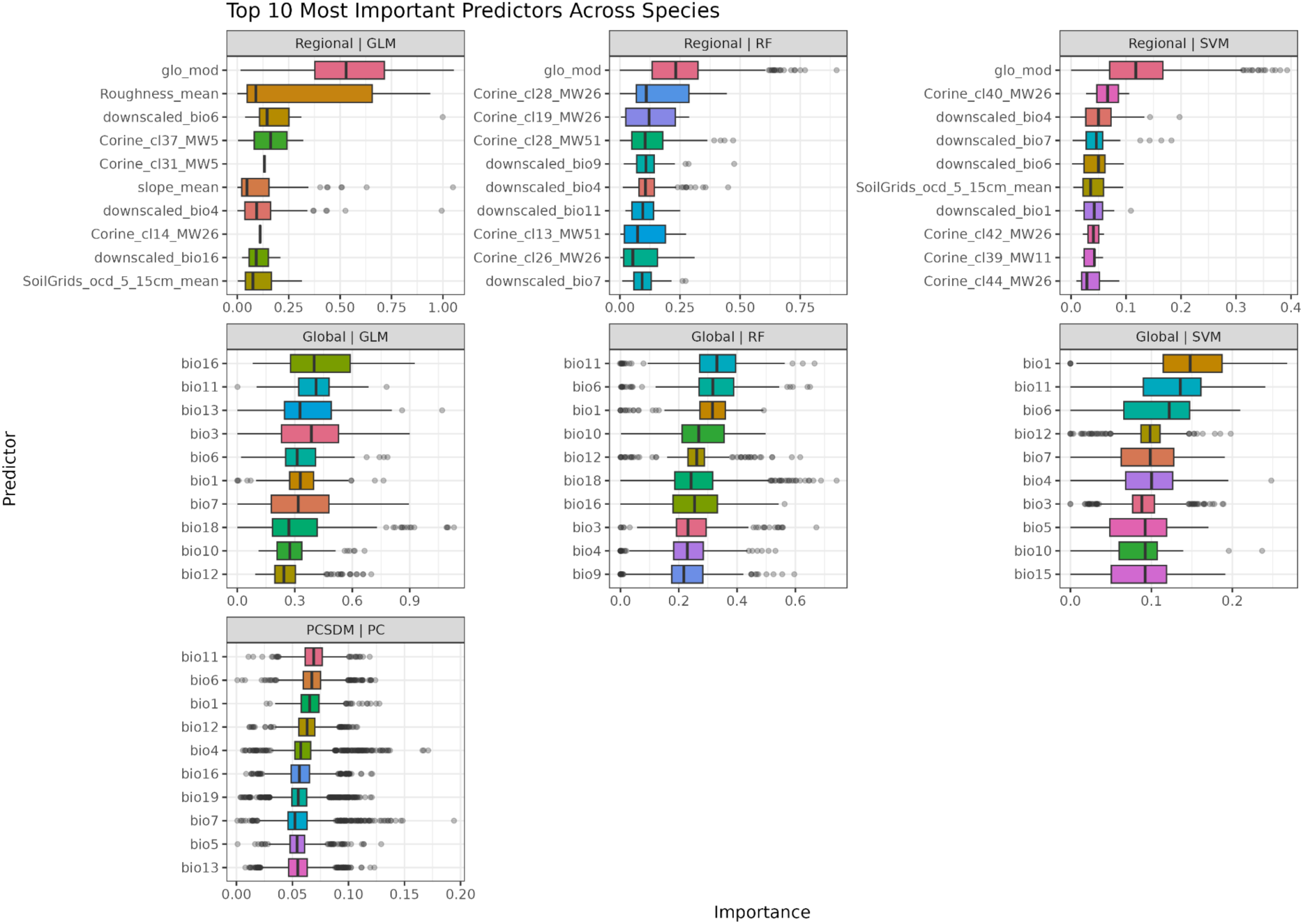
Importance of variables in the species distribution models (SDMS). Only the 10 most important variables for each combination of algorithm and sophistication are shown.

**Figure S4.**
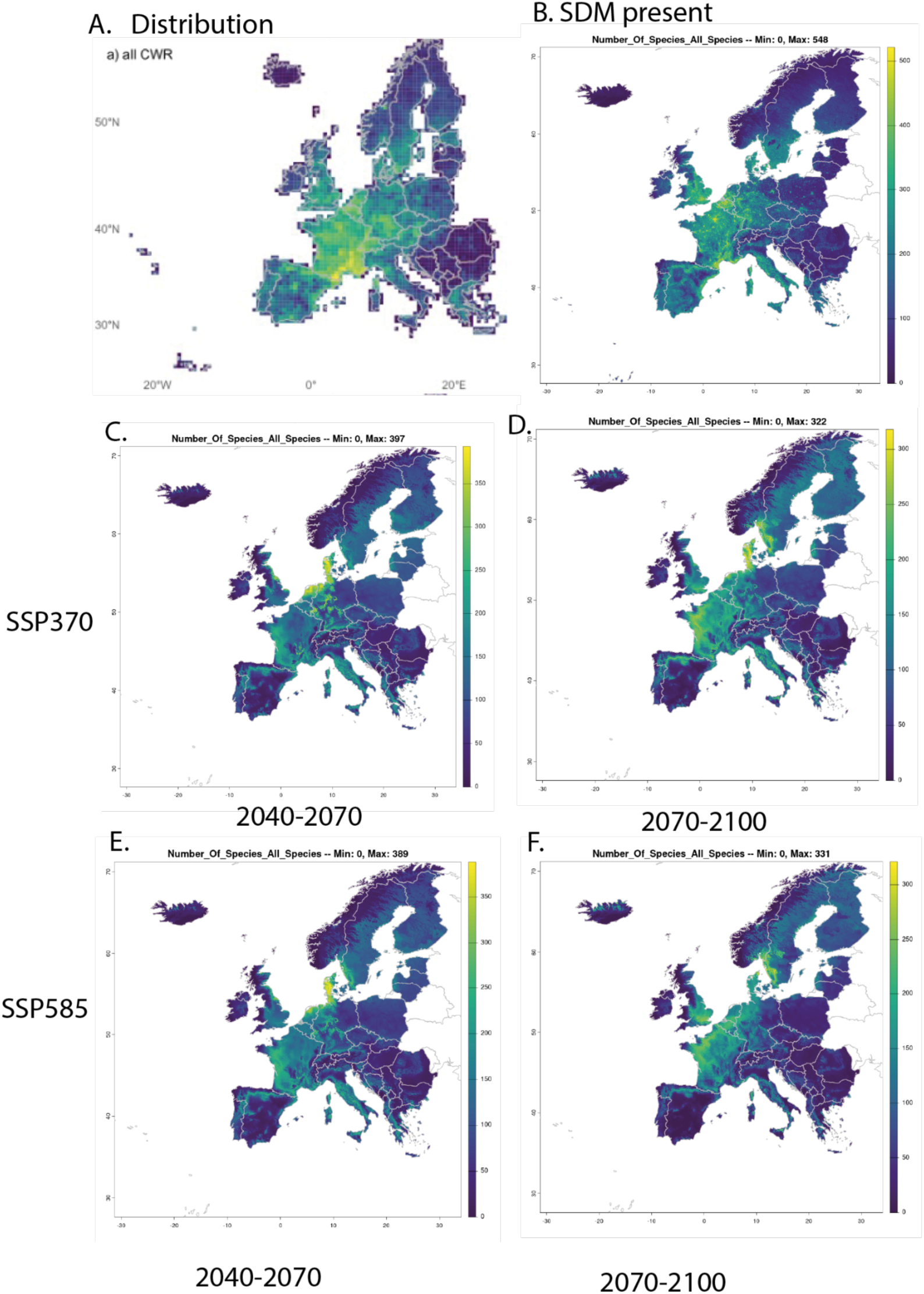
Distribution and diversity pattern of 1140 CWR taxa iin Europe. A. Distribution based on GBIF and Genesys data, resolution: 50km. B. Diversity pattern of current European CWR (resolution: 1 km) and under two different climatic scenarios: SSP370 for C, 2040.2070 and D, 2071-2100 and SSP585 for E, 2040.2070 and F, 2071-2100. Colour scaling is adjusted to the number of species in each condition.

**Figure S5.**
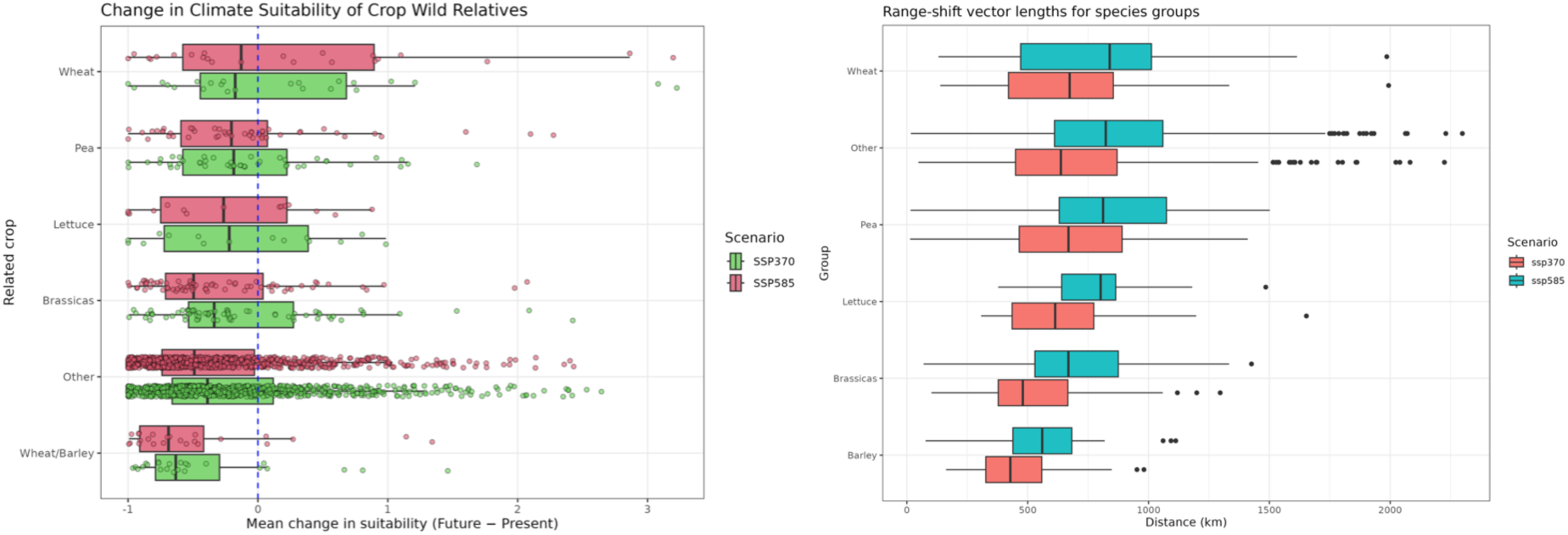
Habitat suitability decline and range-shift distances for European CWRs, grouped by related crop. Boxes show the difference in total predicted habitat suitability between the present and 2071-2100 for SSPs 585 (red) and 370 (green) grouped by parent crop. In spite of several cases of increasing habitat suitability, the average trend for all groups of CWRs is of decline.

**Figure S6:**
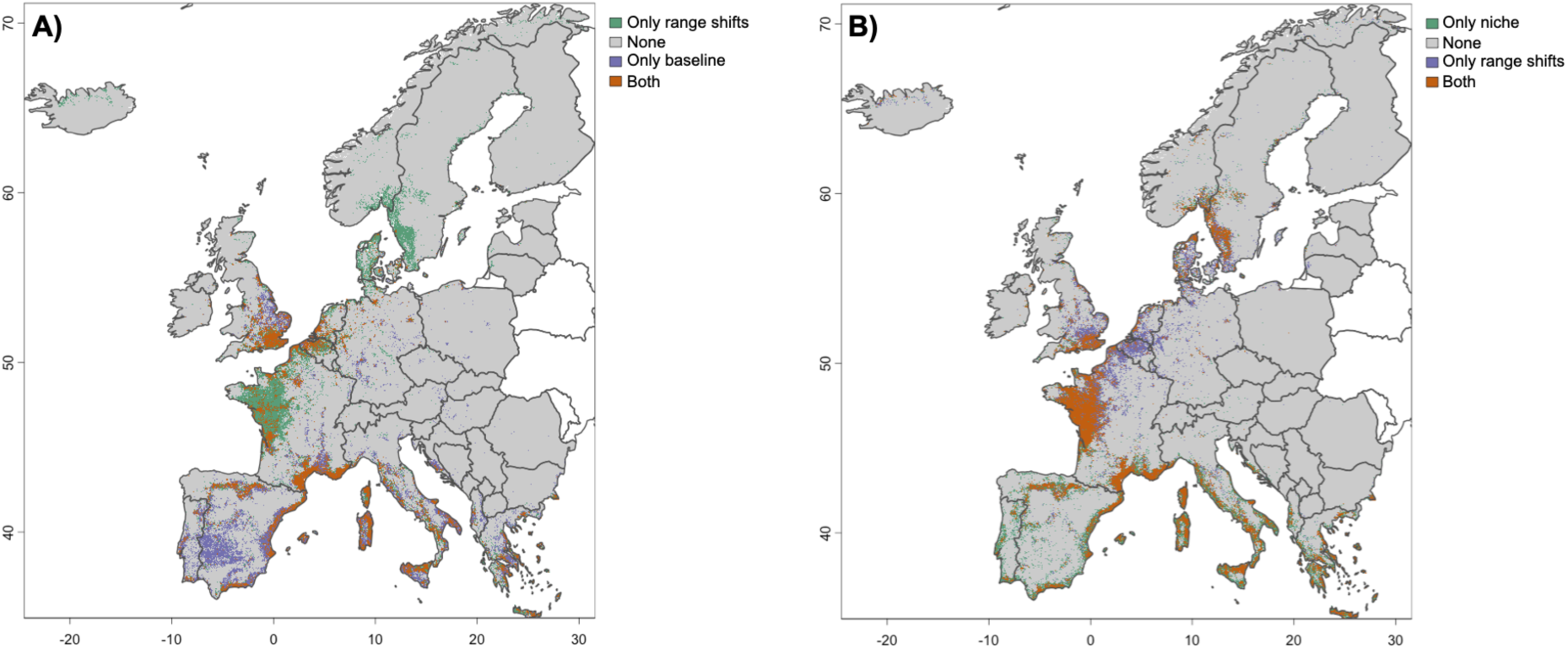
Agreement between baseline, range shifts, and niche models. A: comparison between baseline and range shift. B: comparison between range shifts and niche models.

**Figure S7:**
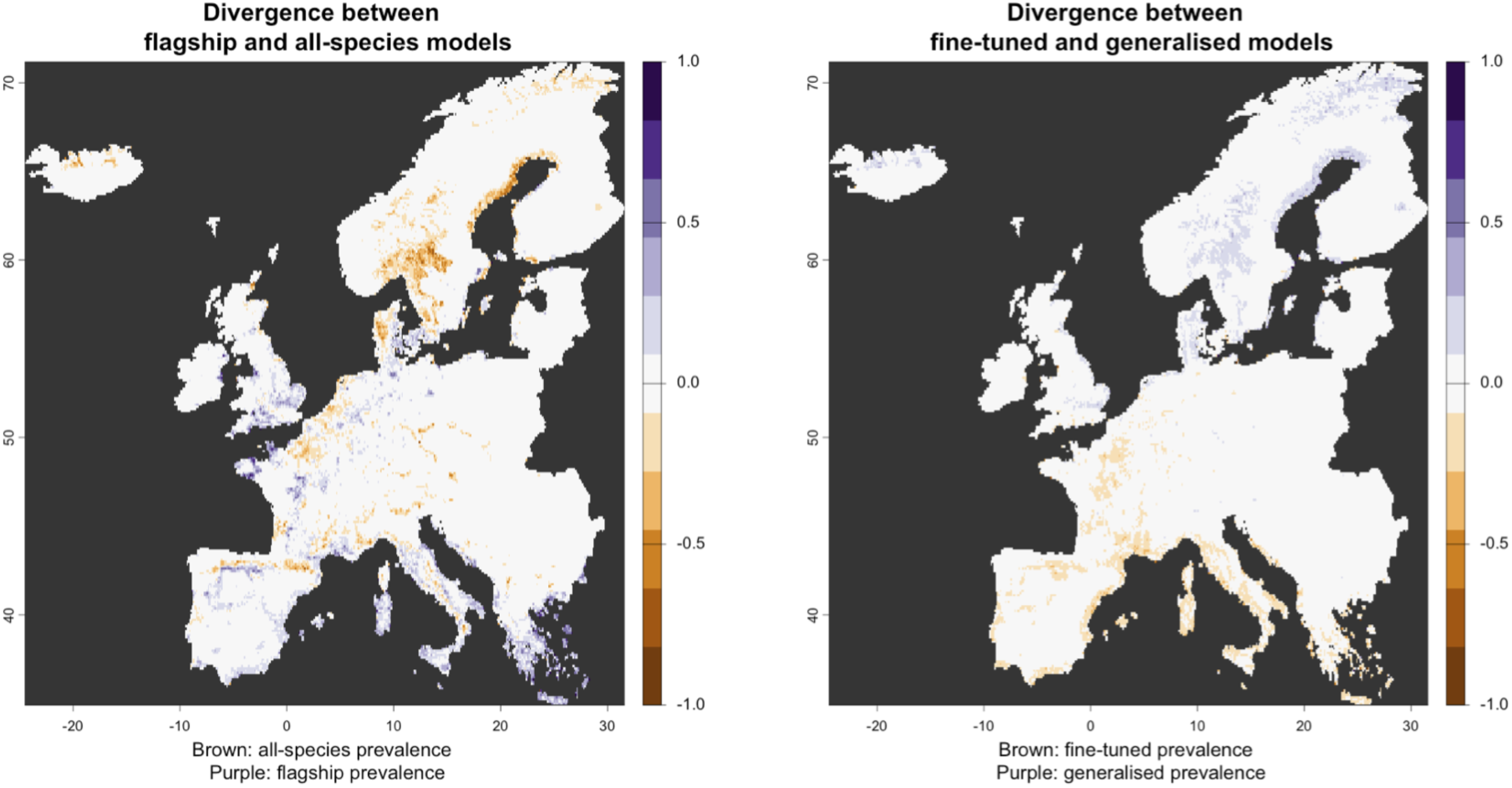
Differences between two pairs of conservation plans. Left: difference between plan accounting for all European CWRs and plan accounting for flagship CWRs only; high-saturation blue and brown areas indicate regions where the flagship CWRs do not act as good umbrella species for the conservation of all European CWRs. Right: difference between two plans, both accounting for all European CWRs, but one trained on all European CWRs (i.e., “fine-tuned” model), the other trained on flagship CWRs only (i.e., “generalised” model); the low divergence values throughout this map indicate only minor differences among these two models, which shows CAPTAIN’s good generalisation performance. To better visualise the fine-scale differences and summarise the level of disagreement between model pairs, we also aggregated these maps to 15 x 15 km^2^ pixels by computing the proportion areas selected only by one model within each 15 x 15 km^2^ cell.

## Supplementary Tables

**Table S1.**
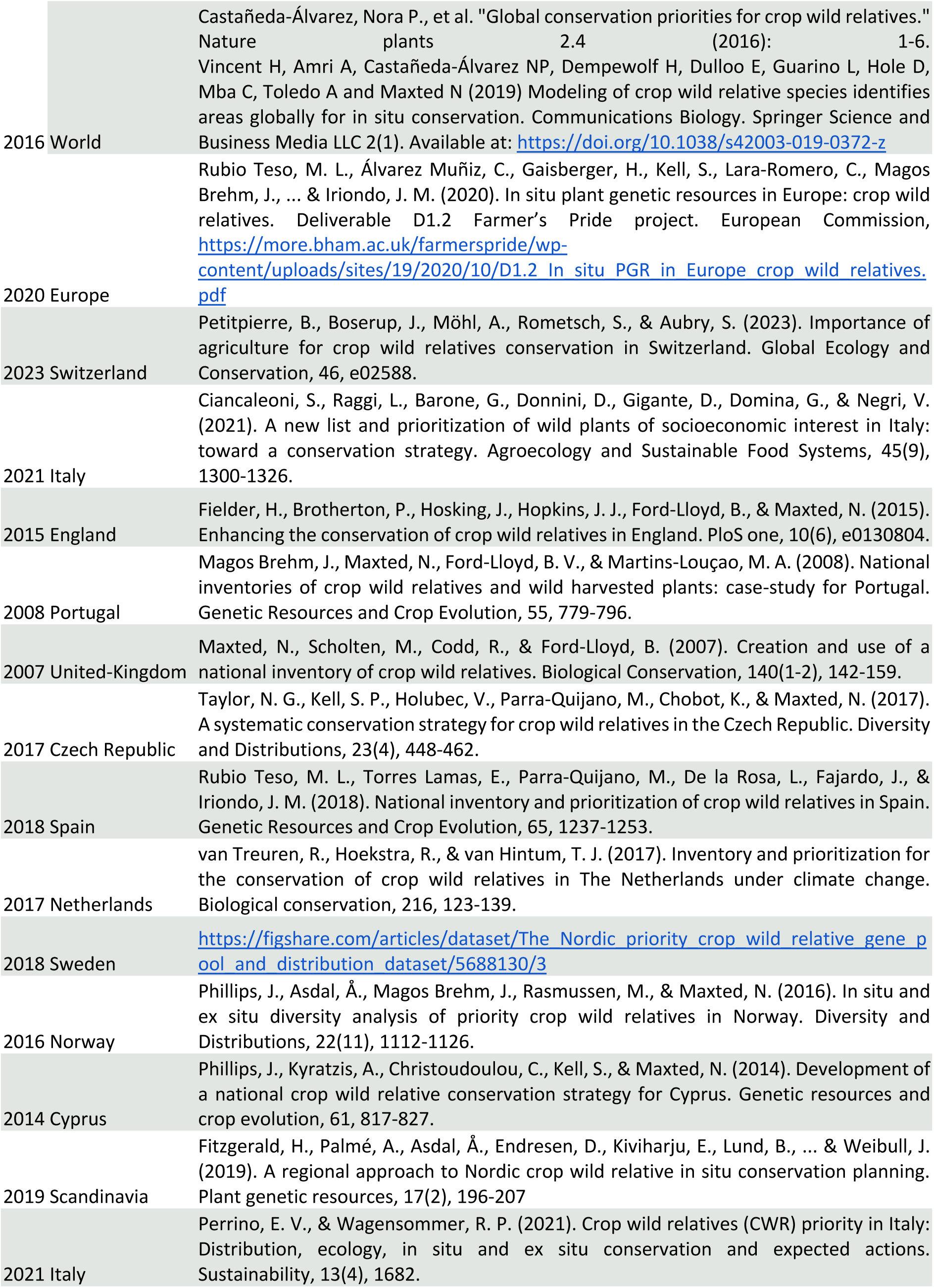
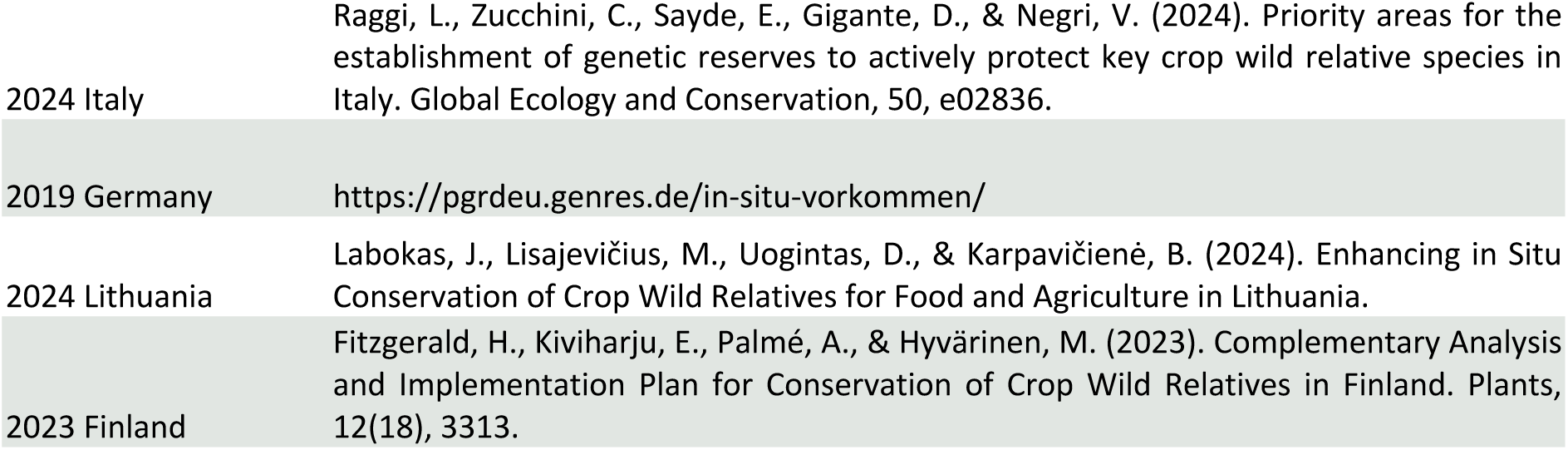
Literature and reports were used to build the European CWR checklist.

|  |  |
| --- | --- |
| 2016 World | Castañeda-Álvarez, Nora P., et al. "Global conservation priorities for crop wild relatives." *Nature plants* 2.4 (2016): 1-6.<br>Vincent H, Amri A, Castañeda-Álvarez NP, Dempewolf H, Dulloo E, Guarino L, Hole D, Mba C, Toledo A and Maxted N (2019) Modeling of crop wild relative species identifies areas globally for in situ conservation. *Communications Biology*. Springer Science and Business Media LLC 2(1). Available at: https://doi.org/10.1038/s42003-019-0372-z |
| 2020 Europe | Rubio Teso, M. L., Álvarez Muñiz, C., Gaisberger, H., Kell, S., Lara-Romero, C., Magos Brehm, J., ... & Iriondo, J. M. (2020). In situ plant genetic resources in Europe: crop wild relatives. Deliverable D1.2 Farmer's Pride project. European Commission, https://more.bham.ac.uk/farmerspride/wp-content/uploads/sites/19/2020/10/D1.2_In_situ_PGR_in_Europe_crop_wild_relatives.pdf |
| 2023 Switzerland | Petitpierre, B., Boserup, J., Möhl, A., Rometsch, S., & Aubry, S. (2023). Importance of agriculture for crop wild relatives conservation in Switzerland. *Global Ecology and Conservation*, 46, e02588. |
| 2021 Italy | Ciancaleoni, S., Raggi, L., Barone, G., Donnini, D., Gigante, D., Domina, G., & Negri, V. (2021). A new list and prioritization of wild plants of socioeconomic interest in Italy: toward a conservation strategy. *Agroecology and Sustainable Food Systems*, 45(9), 1300-1326. |
| 2015 England | Fielder, H., Brotherton, P., Hosking, J., Hopkins, J. J., Ford-Lloyd, B., & Maxted, N. (2015). Enhancing the conservation of crop wild relatives in England. *PloS one*, 10(6), e0130804. |
| 2008 Portugal | Magos Brehm, J., Maxted, N., Ford-Lloyd, B. V., & Martins-Loução, M. A. (2008). National inventories of crop wild relatives and wild harvested plants: case-study for Portugal. *Genetic Resources and Crop Evolution*, 55, 779-796. |
| 2007 United-Kingdom | Maxted, N., Scholten, M., Codd, R., & Ford-Lloyd, B. (2007). Creation and use of a national inventory of crop wild relatives. *Biological Conservation*, 140(1-2), 142-159. |
| 2017 Czech Republic | Taylor, N. G., Kell, S. P., Holubec, V., Parra-Quijano, M., Chobot, K., & Maxted, N. (2017). A systematic conservation strategy for crop wild relatives in the Czech Republic. *Diversity and Distributions*, 23(4), 448-462. |
| 2018 Spain | Rubio Teso, M. L., Torres Lamas, E., Parra-Quijano, M., De la Rosa, L., Fajardo, J., & Iriondo, J. M. (2018). National inventory and prioritization of crop wild relatives in Spain. *Genetic Resources and Crop Evolution*, 65, 1237-1253. |
| 2017 Netherlands | van Treuren, R., Hoekstra, R., & van Hintum, T. J. (2017). Inventory and prioritization for the conservation of crop wild relatives in The Netherlands under climate change. *Biological conservation*, 216, 123-139. |
| 2018 Sweden | https://figshare.com/articles/dataset/The_Nordic_priority_crop_wild_relative_gene_pool_and_distribution_dataset/5688130/3 |
| 2016 Norway | Phillips, J., Asdal, Å., Magos Brehm, J., Rasmussen, M., & Maxted, N. (2016). In situ and ex situ diversity analysis of priority crop wild relatives in Norway. *Diversity and Distributions*, 22(11), 1112-1126. |
| 2014 Cyprus | Phillips, J., Kyratzis, A., Christoudoulou, C., Kell, S., & Maxted, N. (2014). Development of a national crop wild relative conservation strategy for Cyprus. *Genetic resources and crop evolution*, 61, 817-827. |
| 2019 Scandinavia | Fitzgerald, H., Palmé, A., Asdal, Å., Endresen, D., Kiviharju, E., Lund, B., ... & Weibull, J. (2019). A regional approach to Nordic crop wild relative in situ conservation planning. *Plant genetic resources*, 17(2), 196-207 |
| 2021 Italy | Perrino, E. V., & Wagensommer, R. P. (2021). Crop wild relatives (CWR) priority in Italy: Distribution, ecology, in situ and ex situ conservation and expected actions. *Sustainability*, 13(4), 1682. |
| 2024 Italy | Raggi, L., Zucchini, C., Sayde, E., Gigante, D., & Negri, V. (2024). Priority areas for the establishment of genetic reserves to actively protect key crop wild relative species in Italy. *Global Ecology and Conservation*, 50, e02836. |
| 2019 Germany | https://pgrdeu.genres.de/in-situ-vorkommen/ |
| 2024 Lithuania | Labokas, J., Lisajevičius, M., Uogintas, D., & Karpavičienė, B. (2024). Enhancing in Situ Conservation of Crop Wild Relatives for Food and Agriculture in Lithuania. |
| 2023 Finland | Fitzgerald, H., Kiviharju, E., Palmé, A., & Hyvärinen, M. (2023). Complementary Analysis and Implementation Plan for Conservation of Crop Wild Relatives in Finland. *Plants*, 12(18), 3313. |

**Table S2.** Summary Table of the 3 models used.

| Occurrence Threshold | Model Name | Predictor Variables | Calibration | Prediction | Methods | Nb. CWRs |
| --- | --- | --- | --- | --- | --- | --- |
| More than 5 occurrences | PC Model | 19 Bioclim | World | Europe | Niche projection | 1140 |
| More than 30 occurrences globally | Global model | 19 Bioclim | World | Europe | Ensemble of GLM, RF, SVM | 915 |
| More than 30 occurrences in Europe | Nested European model | 19 Bioclim; microclimate; land-use; soil; topography<br>Global model | Europe | Europe | Ensemble of GLM, RF, SVM | 819 |

**Table S3.** Summary of the variable classes used in the three levels of model sophistication.

| **Sophistication** | **Variables class** | **Source** |
| --- | --- | --- |
| Global | Chelsa V 2.1. BIOCLIM variables set | https://www.chelsa-climate.org/models/chelsa |
| PCM | Chelsa V 2.1. BIOCLIM variables set | https://www.chelsa-climate.org/models/chelsa |
| Regional | Chelsa V 2.1. BIOCLIM corrected for canopy shading based on ForestTemp (Haesen et al., 2021) | Flavien Collart, personal communication. |
| Regional | Corine land-use at multiple moving windows | https://land.copernicus.eu/en/products/corine-land-cover |
| Regional | Terrain roughness | Copernicus' 25 m x 25 m Digital Elevation Model |
| Regional | Soil chemical physical properties | https://doi.org/10.5194/soil-7-217-2021 |
| Regional | Solar radiation | SI, Solar radiation |
| Regional | Soil wetness index | SI, Wetness index |

**Table S4:** Interpretation of AUC values and TSS values (according to Allouche et al. 2006) produced for each species and each modelling technique based on cross-validation.

| **AUC** | **Model quality** | **TSS** | **Model quality** |
| --- | --- | --- | --- |
| > 0.9 | Excellent | > 0.8 | Excellent |
| 0.8–0.9 | Good | 0.6–0.8 | Good |
| 0.7–0.8 | Fair | 0.4–0.6 | Fair |
| 0.6–0.7 | Weak | 0.2–0.4 | Weak |
| <0.6 | Fail | <0.2 | Fail |

